# Does plant diversity determine the diversity of the rhizosphere microbial community?

**DOI:** 10.1101/2021.08.18.456889

**Authors:** Aleksei Zverev, Arina Kichko, Vasiliy Shapkin, Aleksandr Pinaev, Nikolay Provorov, Evgeny Andronov

## Abstract

The rhizosphere community represents an “ecological interface” between plant and soil, providing the plant with a number of advantages. Close connection and mutual influence in this communication allow to talk about the self-adjusting “plant-rhizosphere community” system, which should be be studied in connection. Diversity estimation is one of the ways of describing both bacterial and plant communities. Based on the literature, there are two assumptions of how the diversity of plant communities related to the diversity of bacterial communities: 1) an increase in the species richness of plants leads to an increase in the number of available micro-niches, and increasing of microbial diversity, 2) an increase in the species richness of plants is accompanied by the predominant development of bacteria from highly productive specific taxa and decreasing in the diversity of microorganisms. Experimental studies show controversial results.

We analyzed field sites (rye crop field and two fallow sites), using DNA isolation of both the plant root mass (followed by sequencing of the ITS1 region) and rhizosphere microorganisms (followed by sequencing of the 16s rDNA V4 region). This allowed us to 1) accurately determine the abundance and taxonomic position of plant communities; 2) extract information about both plant and microbial communities from the same sample.

There was no correlation between alpha-diversity indices of plants and rhizosphere communities. Alpha-diversity connection should be explored in similar plant communities, such as synusia. We hypothesize, that the significant differences in plant abundances lead to significant changes in exudation profiles, and the loss of diversity connection. The beta-diversity between rhizosphere communities and plant communities is highly correlated, in particular in terms of the abundance of taxa. This can be explained by a potential correlation (as reported in the literature) or by the presence of statistical artifacts.

## Introduction

As the formation of a specific microbial community near the plant root, the phenomenon of the rhizosphere effect has been the subject of many works of both classical and modern biology. The rhizosphere community represents an “ecological interface” between plant and soil, providing the plant with a number of advantages such as growth stimulation, protection from pathogens, nutrition, among others [1, 2]. The source of the rhizosphere microbiome is both the microbial community of plant seeds [3] and the community of soil microorganisms [4]. The composition and abundance of plant root exudates determine the formation of the bacterial community [5, 6]. The source of the microbiome, and the development of it, thus forms the final community.

Diversity in both sources and ways of development lead to the specificity of the rhizosphere microbiome. Several factors can affect the composition of rhizosphere communities. In addition to the type and agrochemical properties of the soil, the genotype of the plant (species [7, 8] and cultivar [9]) is a significant factor for microbiome development. This can be explained by the specific exudation spectrum from various plants, as the spectrum of secreted substances depends not only on the species or cultivar but also on the developmental phase, physiological state, etc. [10, 11]. In turn, the microbial communities themselves affect the metabolic status of the plant [12], which allows us to talk about the “plant-rhizosphere community” system as a self-adjusting system (a kind of “gut-brain axis” in plants). This system is additionally complicated by the high diversity of plant species, which is natural in indigenous plants populations.

Diversity estimation is one of the ways of describing both bacterial and plant communities. As self-adjusting systems, rhizosphere microbiomes and plant communities are connected, and therefore, the following question arises: is the diversity of plant communities related to the diversity of bacterial communities? Based on the literature, there are two assumptions: 1) an increase in the species richness of plants leads to an increase in the number of available micro-niches, which leads to an increase in microbial diversity, 2) an increase in the species richness of plants is accompanied by the predominant development of bacteria from highly productive specific taxa, which leads to a general decrease in the diversity of microorganisms [13]. Experimental studies of this relationship show controversial results: some studies indicate the absence or negative correlation between plant and bacterial richness [14, 15], whereas others show the presence of a positive relationship [16]. Beta diversity indices show an unambiguous positive correlation between the distances of plant and bacterial communities [17].

There are examples of research of similar relationships between the diversity of plants and the associated microorganisms. A close relationship was demonstrated in a study of the diversity of the rhizobial soil community (by the nodA gene) and the diversity of their symbiotic hosts (the NFR5 and K1 genes of leguminous plants), allowing the authors to formulate the hypothesis of “evolutionary molding”, where the plant community plays the role of the rigid matrix and the microbial community acts as a “molding” substance. The whole process not only links diversities, but even phylogenetic topologies; for the description of the last phenomenon, the concept of beta-topological diversity has been introduced [18]. Another example is the close relationship between the diversity of *Galega* rhizobia and their two hosts, *G. orientalis* and *G. officinales*, revealed at the level of genomic AFLP fingerprints [19]. However, for less closely integrated systems with indigenous plant communities, analyzed by taxonomic rather than functional markers, such a relationship has not yet been shown.

One of the reasons for the uncertainty in this area is, most likely, the problem of the correct estimation of plant diversity, for which the same algorithms that are used today for the analysis of microbial diversity could be applied. The development of NGS methods for the estimation of microbial diversity, from DNA extraction methods to statistical analysis of libraries (16S, ITS), has led to the rise of this area observed today [20]. However, in most papers, plant diversity is determined using geobotanical methods [13, 15], but this approach is not accurate enough in both determining the species in the communities and their abundances [21]. In addition, in the context of the aim of this study, there is a question about the correspondence between the “aboveground” and “belowground” plant diversities, which can be quite different [22].

Therefore, in this work, we analyzed the diversity of the plant population via direct DNA isolation of the plant root mass, followed by NGS sequencing of the ITS1 region; rhizosphere soil samples were taken from the same root sample. This allowed us to 1) accurately determine the abundance and taxonomic position of plant communities; 2) extract information about both plant and microbial communities from the same sample; 2) analyze plant diversity using the same NGS approaches as used for the rhizosphere community of microorganisms. Thus, we were able to use the same algorithms and metrics to analyze both communities.

## Materials and Methods

Samples were collected on July 21, 2017, on fields of the Pskov Research Institute of Agricultural Sciences and Rodina State Farm in the Pskov region, Russia (coordinates of the collection point are 57.845611 N 28.201028 E). We select one site within a rye crop field (referred to as Monoculture Rye or MonoR) and two fallow sites outside the field border from two locations, dominated by cereals (Polyculture Cereal or PolyC) and *Galium* and *Dactylis* species (Polyculture Galium, or PolyG). In each site, three samples were taken. The photo and the geobotanical description of the sampling sites is provided in the S1 Table and S1 Fig.

Bricks of topsoil (about 15 × 15 × 10 cm) were collected and stored in individual packages not longer than 48 hours. In the laboratory, bulk soil was gently removed manually by shaking, and 30 g of the root mass was intensively shaken with 50 ml of 0.005M Na-phosphate buffer in a Pulsifier II (Microgen, UK) in provided bags (PUL512 Bags) for 1 min. The liquid fraction was centrifuged, and the pellet was used to isolate the total rhizosphere DNA. The root mass was used to isolate plant DNA.

Procaryotic DNA from the pellet was isolated using the MN NucleoSpin Soil Kit (Macherey-Nagel, Germany) and a Precellus 24 homogenizer (Bertin, USA). Quality control was carried out by PCR and agarose gel electrophoresis. Sequencing of the V4 variable region of the 16S rRNA gene was performed on an Illumina MiSEQ sequencer, using the primers F515 (GTGCCAGCMGCCGCGGTAA) and R806 (GGACTACVSGGGTATCTAAT) [23].

Plant DNA from roots was isolated using mechanical destruction in liquid nitrogen, followed by phenol extraction; the quality of the DNA was also checked via agarose gel electrophoresis. Sequencing of the ITS1 variable region was performed on an Illumina MiSEQ sequencer, using the primers ITS-p5 (YGACTCTCGGCAACGGATA) and ITS-u2 (GCGTTCAAAGAYTCGATGRTTC) [24].

The general processing of sequences was carried out in R 3.6.4, using the dada2 (v. 1.14.1) [25] and phyloseq (v. 1.30.0) [26]packages. For taxonomic annotation, the databases SILVA 138 [27] and PLANiTS [28] were used.

The main alpha- and beta-metrics were calculated using the phyloseq and picante [29] packages. For the mean p-distance in a library, we used the home-brew script with following steps: 1) make multiple alignment for reference sequences; 2) extract p-value for every pair of sequence; 3) multiple this p-value to abundance of both seqences; 3) sum all values. Correlations between diversity indices were calculated using Spearman correlation. Significant differences in abundances of taxa between sites were determined using theDeSEQ2 package [30].

All reads were submitted to SRA (PRJNA649486) and are available under the link https://www.ncbi.nlm.nih.gov/sra/PRJNA649486.

## Results

The ITS1 sequencing of plant DNA yielded 230 ASVs. The taxonomic position at genus level was defined for 217 of them; the number of reads per sample after rarefaction was 14,210. Regarding 16s rDNA, we found 5,284 ASVs, with 15,487 reads per sample.

Fig 1 shows the taxonomic composition of the communities. The geobotanic description (provided in the S1 Table) corresponded with the composition structure according to ITS1 sequencing (Fig 1A). Almost all reads from MonoR libraries were attributed as *Secale;* PolyC (samples from the cereal synusia) had about half the reads from *Poa*, followed by *Elymus* and *Dactilus*. PolyG (samples from *Galium* and *Dactylis* synusia) was more diverse; most reads were attributed as *Galium, Poa*. Despite the geobotanical description of this synusia as mixed *Galium* and *Dactylis*, there was no evidence of a great amount of *Dactylis* in the PolyG libraries, which can be explained by difficulties in describing the graminae vegetation outside the flowering phase.

**Fig 1.**
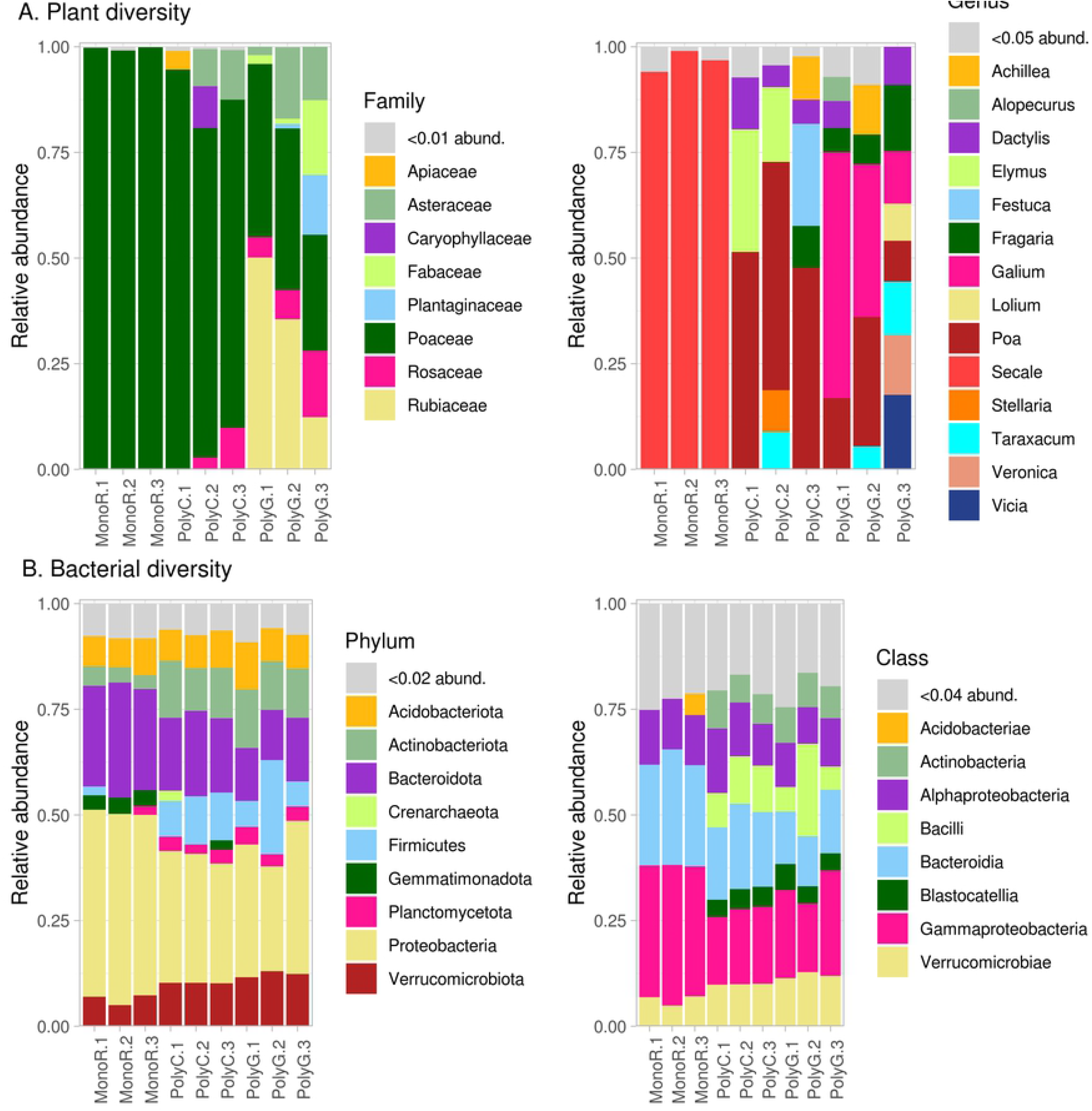
Relative abundances of plant and bacterial communities in the experimental fields.

The bacterial communities from the samples (Fig 1B) were typical for rhizosphere microbiomes [11]. The communities from the two fallow sites (PolyC, PolyG) were similar, whereas the communities from the rye site (MonoR) contained more *Gammaproteobacteria* (*Proteobacteria*) and *Bacteroidia* (*Bacteroideta*). Differential abundance analysis allowed us to find 584 microbial taxa, which significantly differed between two sites, with information about abundance in repeats. When comparing PolyC and PolyG, none of the OTUs was marked as significantly different. The results of the pairwise comparison of PolyC and PolyG with MonoR are shown in Fig 2 at the order level.

**Fig 2.**
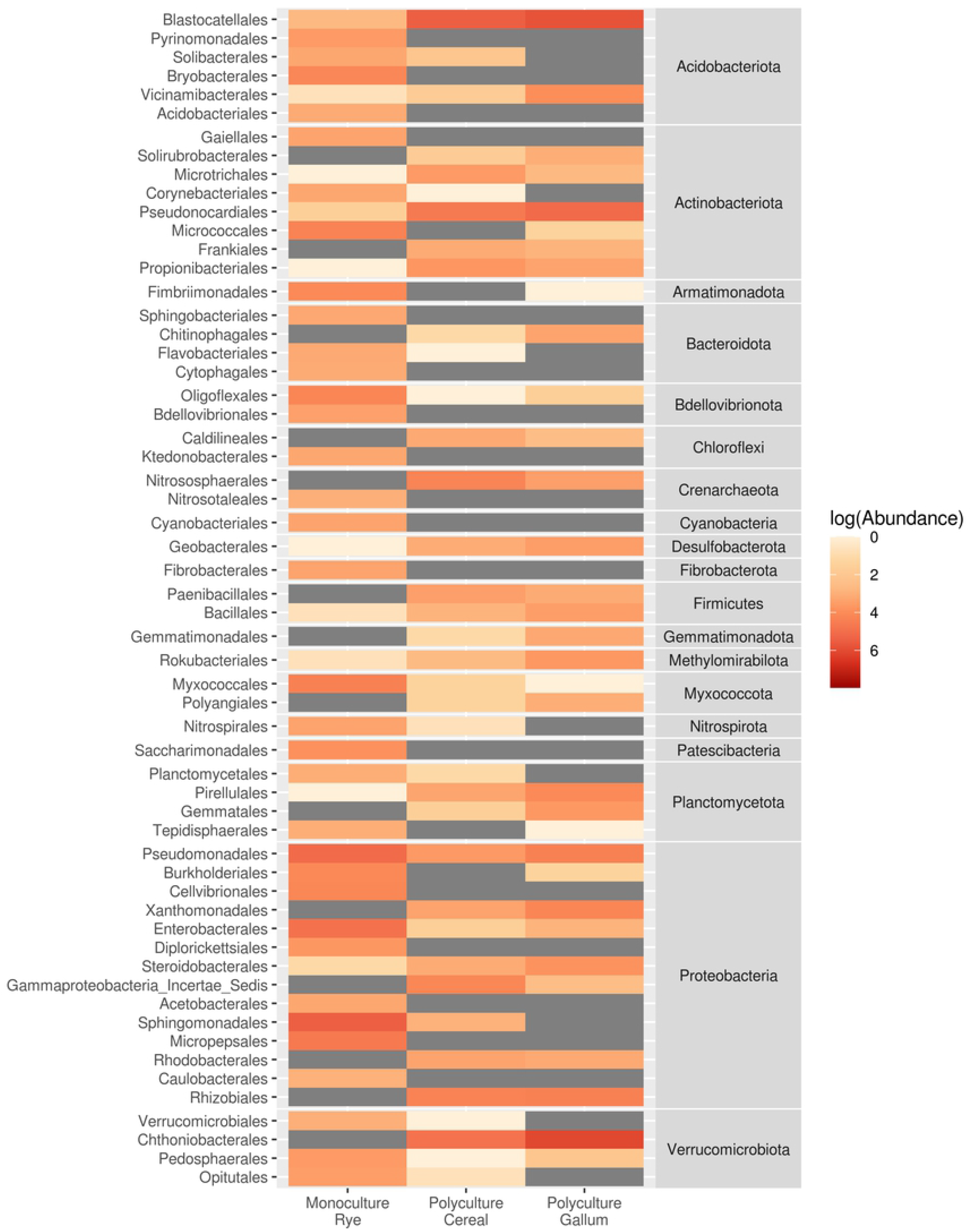
Mean abundances of bacterial orders with different inter-source abundances (according to DeSEQ2)

In this comparison, the most abundant OTUs in rye crop rhizosphere microbiomes were from *Proteobacteria* (orders *Pseudomonadales, Sphingomonadales, Enterobacteriales*), followed by *Armatimonadota* (*Fimbriimonadales*), *Acidobacteriales* (*Bryobacterales*), and *Actinobacteriota* (*Micrococcales*). In fallow root microbiomes, *Verrucomicrobia* (*Chthoniobacteriales*), *Acidobacteria* (*Blastocatellales*), and *Actinobacteria* (*Pseudonocardiales*) were more abundant. For all samples, the most common alpha-diversity indices, observed OTUs, Shannon, Simpson, mpd (mean pairwise distance), and p-dist (mean p-distance, restored from alignment, see Materials and Methods), were calculated at different taxonomic levels (Fig 3).

**Fig 3.**
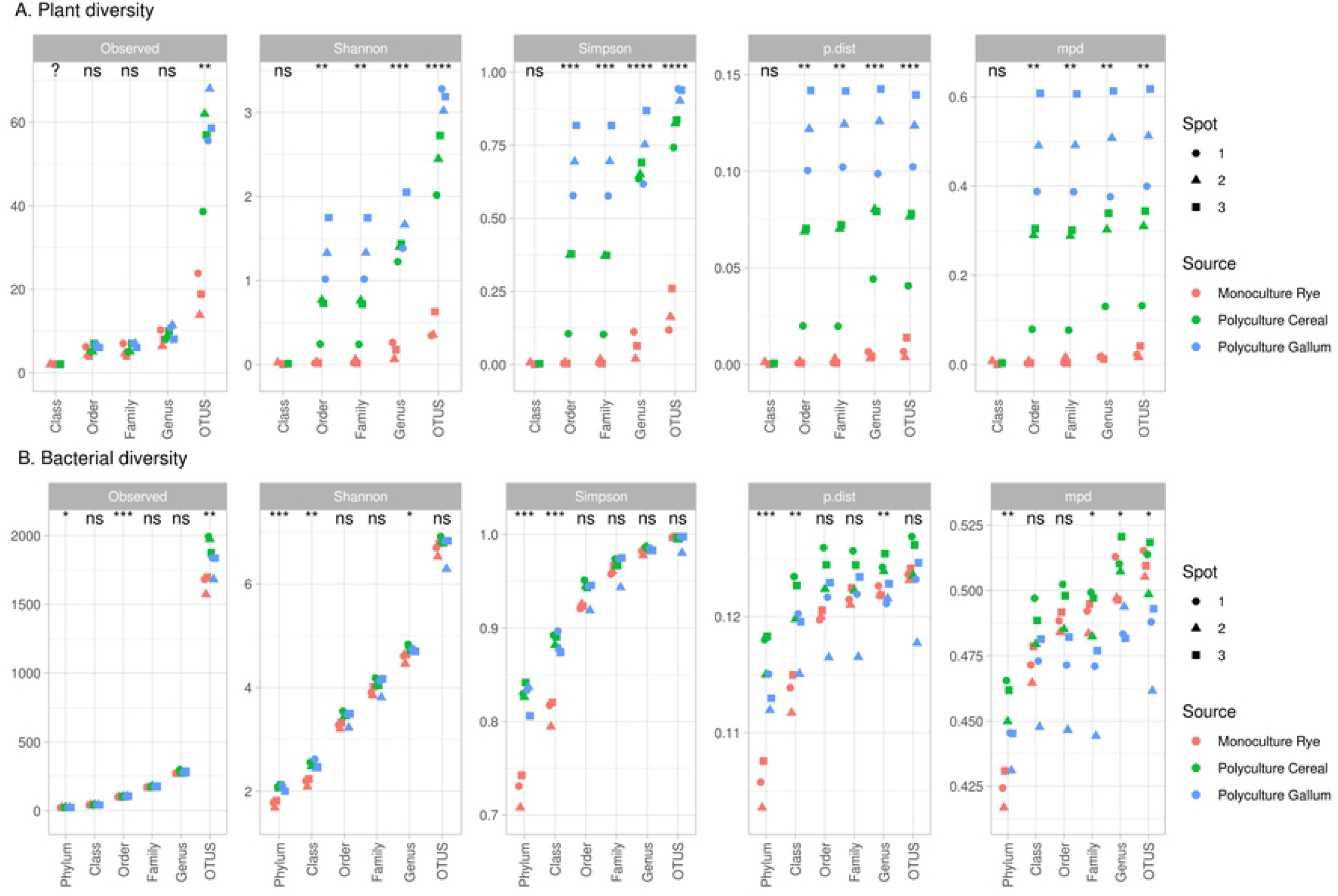
Alpha-diversity indices for plant and bacterial communities (significance for ANOVA in groups; ns – non-significant, * - p < 0.05, ** p < 0.01, *** p < 0.005)

The plant communities were highly diverse at different levels and indices (Fig 3A). The observed OTU index was not useful because of the presence of weedy plants in the rye crop. Weighted indices, such as mpd or p-dist, were more suitable; the differences between sample sites were significant for both indices up to the order level.

The bacterial communities were less diverse. The observed OTUs as well as the p-dist and mpd indices showed that communities from the PolyC site were more diverse, mostly at low taxonomical levels (OTUs, genus). Interestingly, samples from rye roots (MonoR) were significantly less diverse at phylum level, whereas at other taxonomical ranks, there were no such differences.

For each index on the OTU taxonomic level, the correlation between plant diversity indices and microbial diversity indices was calculated. We observed no significant correlations between plant and rhizosphere microbial diversity.

Fig 4 shows the beta-diversity indices (Bray distance and weighted UniFrac). According to the weighted UniFrac index (Dunn index for roots 0.316, for microorganisms 0.821), the samples were mixed (Fig 4A). Samples from rhizosphere communities from PolyC and PolyG were mixed in one cluster. While the MonoR and PolyC samples were close, the PolyG samples were different. This might be connected to the phylogenetic composition of the population; *Secale* and *Poa*, the two main genera from these sites, both belong to the *Poaceae* family, whereas *Galium* belongs to the Rubiaceae family (Fig 1A).

**Fig 4.**
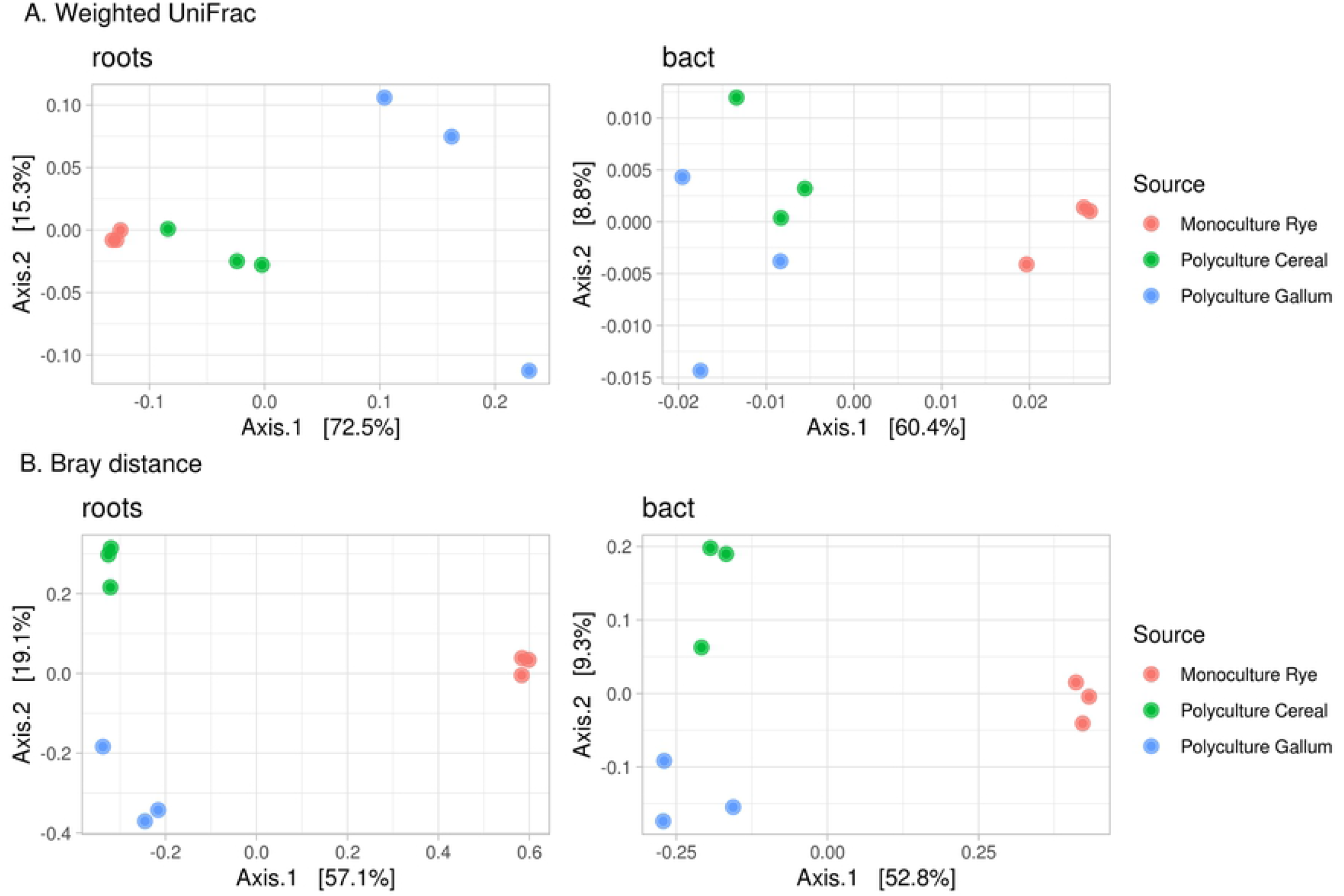
Beta-diversity indices for plant and bacterial communities.

According to the Bray distance (Dunn index for roots 0.832, for microorganisms 0.909), all communities, from plants and from rhizospheres, formed their own clusters (Fig 3B).

The correlation between the distance matrix in plant and bacterial communities was high for the Bray distance (R = 0.866, p = 0.01) and not significant for the weighted UniFrac index. Fig 5 shows the results of the beta-distance correlation, excluding the intra-cluster distance (distance between repeats of the same sample).

**Fig 5.**
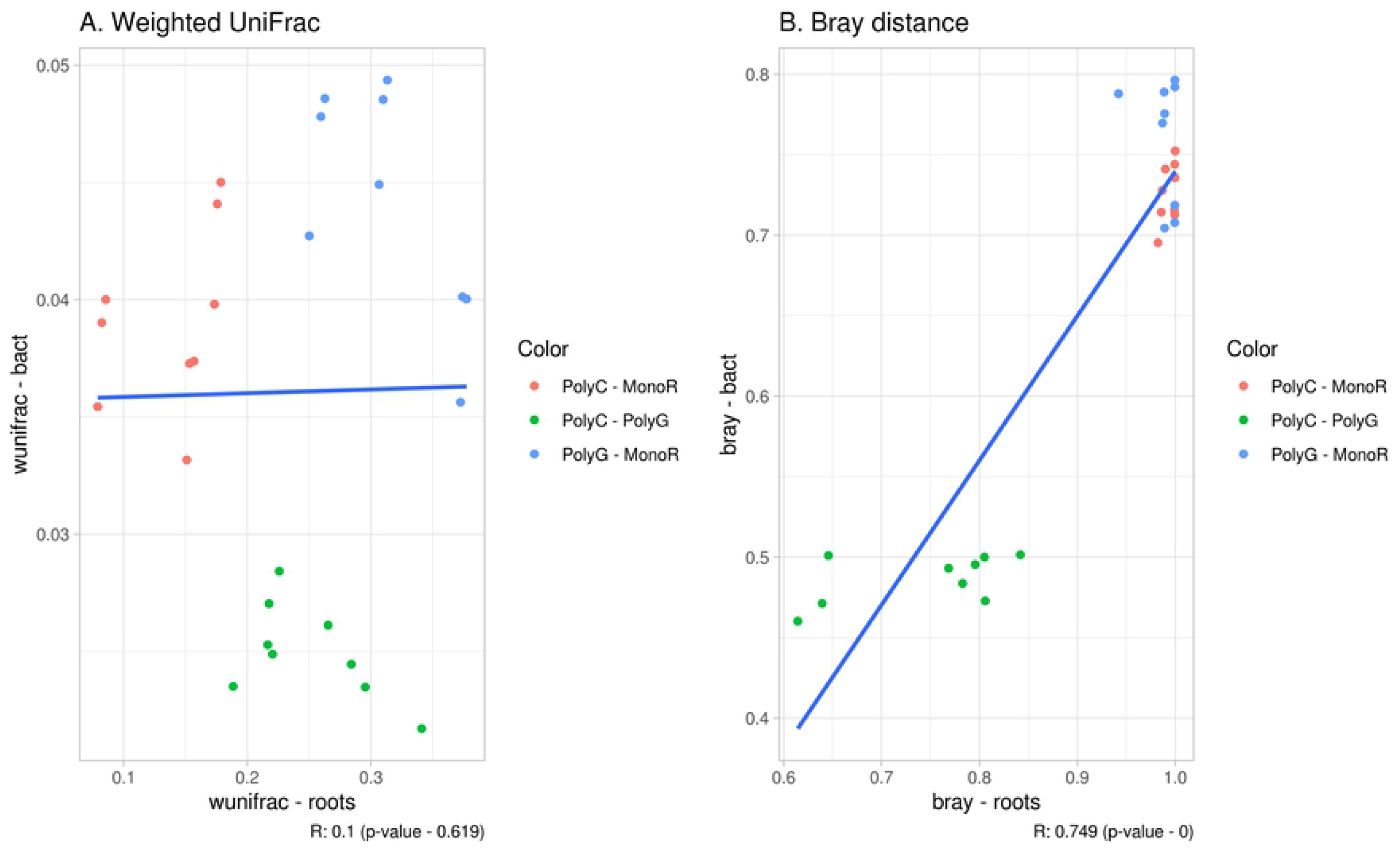
Correlation between beta-diversity indices.

## Discussion

The aim of this work was to determine the potential connection between the diversity of plant communities and the diversity of rhizosphere microbiomes. For this purpose, we used three sites with different plant communities (synusiae) in the same location: a rye crop monoculture (MonoR) and two polyculture fallow sites, dominated by cereals (PolyC) and *Galium* and *Dactylis* (PolyG). Despite differences in crop species and tillage, soil type, water regime, and main soil properties were similar for all three sites, minimizing the influence of these factors on microbial communities.

The novel approach to estimate plant community composition (ITS1 sequencing) is highly effective. Almost all (217 of 230) determined ASVs were annotated, and the composition and abundance of plant genera fit the geobotanical description. In addition, this method allows 1) to characterize plant communities via their underground parts (which interact with rhizosphere microorganisms); 2) use the same sample for both plant and microbiome analysis (therefore, each rhizosphere library matches with the plant library); 3) use standard approaches for processing and calculating diversity indices and generating bar graphs.

For samples obtained from same soil and location, the most important factor shaping the microbial community of the rhizosphere is the plant community structure. We hypothesize that the variation in the spectrum of exudates could be one of the possible mechanisms responsible for the diversity of the rhizosphere microbial community. This spectrum is specific for various plant species and genotypes, and therefore, in the case of the whole plant community, we can talk about an “exudome” of the community, whose structure depends on the abundance of plants in the community. This exudome can be described by the number of the individual plant spectra and by their weights (amount of exudate spectra of exact plant species in all exxudome), according plant abundances. In the comparison of two polycultures, the number of plant species is the same (as shown in Fig 3A), with variations in abundance; it therefore seems logical that the exudomes of these communities also will differ not in the number of different spectra but in their weights. In contrast, when comparing monoculture plants communities with polyculture ones, there are differences in the number of plant species. In this case, exudomes of plant communities will differ in the number of different spectra and in their weights.

In this study, using modern approaches, we found no correlation between different alpha-diversity indices. According to all indices, rhizosphere communities in the monoculture site (MonoR) were as diverse as those in the polycultures (PolyR, PolyC), whereas the plant community diversity in the monoculture was obviously lower. Regarding the polyculture (and mpd and p-dist indices), more diverse plant communities (PolyG) showed a less diverse rhizosphere microbial community. This might be evidence of a negative connection, as suggested by Goberna and coauthors [13], but this connection can be found only in plant communities with close exudomes (differing in abundance by not in the number of spectra). Significant differences in exudomes can lead to different microbial response and a loss of similarity, when environmental factors override the realistic relationship. Perhaps, this variation in exudome abundance allowed Schlatter and colleagues to show that in artificial plant communities, there is a decrease in bacterial diversity with increasing richness (1, 4, 8, 16 species) [31].

Despite the expected main difference in plant diversity between poly- and monocultures, with a moderate variation between polycultures, both alpha- and beta-diversity of the plants showed significant variations between synusiae. In the community of different cereals, PolyG in most cases is significantly more diverse by all alpha-diversity indices, and shows inter-sample variability (sample PolyG.3). This effect is also clear at higher taxonomic levels (family or order).

Differential abundance analysis did not show any difference between rhizosphere communities of polycultures. Similarly, OTUs and Shannon indices of these samples were also not significantly different. Only phylogenetic-related indices (mpd and p-dist) were significantly different, leading us to infer that the phylogenetic composition of communities varies in the rhizospheres of similar plant communities. However, further elaboration of this theory is recommended.

In contrary, when comparing the microbial communities of the polyculture sites with those of the monoculture, there was a significant difference in the observed OTUs, whereas phylogenetic-related indices (mpd, p-dist) showed no clear pattern. Differential abundance analysis enabled us to identify taxa with significant differences in relative diversity, most likely because of significant differences in exudome profiles and the loss of similarity between plant-microbial systems.

This idea also fits well with the beta diversity plots. Indeed, the distances between samples from polycultures were small for both plant and bacterial communities (about 17% of the explained variance in PcoA plots for plants and 8% for microorganisms). The distances between samples from mono- and polycultures were large (about 60% of the explained variance in PcoA plots). This allowed us to estimate a correlation for Bray distances only (due to the taxonomic similarity between rye and cereal, the correlations between weighted Unifac distances were not significant). This correlation might be a true correlation, as observed in previous studies [17], or a statistical artifact (caused by the large distance between monoculture and polyculture clusters).

The significant distance in beta-diversity is connected to the differences in the abundances of the different taxa. Hypothesized as a molding matrix, differences in abundances between plant communities are not useful, whereas abundances of microbial taxa represent the specificity of rhizosphere taxa in different plant communities (at the order level, as shown in Fig 2). Some of them have previously been reported as rhizosphere taxa. *Sphingomonadales* and *Enterobacteriales* are typical for plant rhizosphere communities [32], and *Fimbriimonadales* has previously been described as a bacterial taxon from the *Anthurium andraeanum* rhizosphere community [33]. In the same case, *Blastocatellales* has previously been described as an oligotrophic, slightly acidophilic to neutrophilic mesophile from arid soils [34]. However, these data are controversial, as they can describe a novel location of this microorganism or a biased database. More precise data can be obtained by functional or full-genomic analyses of rhizosphere communities.

Despite the absence of a clear correlation between plant and bacterial diversity, comparing the results with the previously mentioned concept of evolutionary molding, where the plant population “molds” in the microbial population and where the correlation between plant and microbial diversities is clear, is challenging. The reasons are as follows: 1) the symbiotic system itself is tighter than the rhizosphere community interaction; 2) in this work, we used regular taxonomic marker genes, not specific plant or bacterial genes. A closer interaction could be revealed with the use of genes directly involved in plant-rhizosphere communication processes. This could be plant genes involved in plant root exudation and responses to bacterial signals (for example, strigolactone-processing genes) and bacterial genes involved in the decomposition of those exudates by microorganisms. Also, it is important to characterize the structure and composition of common exudation profiles of different plant communities.

## Conclusions

1. Sequencing of the ITS1 region is highly effective for the taxonomical annotation of plant communities. Almost all ASVs were attributed; abundances of taxa fairly corresponded to the geobotanical descriptions.
2. There was no correlation between alpha-diversity indices of plants and rhizosphere communities. Alpha-diversity connection should be explored in similar plant communities, such as synusia. Significant differences in plant abundances lead to significant changes in exudation profiles, different microbial responses, and the loss of diversity connection.
3. In contrast, the beta-diversity between rhizosphere communities and plant communities is highly correlated, in particular in terms of the abundance of taxa. This can be explained by a potential correlation (as reported in the literature) or by the presence of statistical artifacts.
4. In future studies, the diversity connection should be analyzed by searching for functionally related genetic regions of plants and soil microorganisms.

## Acknowledgments

This work was performed using the equipment of the Genome Technologies and Cell Biology Common Use Center, All-Russian Research Institute of Agricultural Microbiology.

## Supporting information

**S1 Fig. General view, current location and root samples**. MonoR — rye crop field; PolyC - forb and cereal meadow dominated by cereals ; PolyG - forb and cereal meadow dominated by *Galium* and *Dactylis* species. Samples are: 1 — MonoR.1; 2 — MonoR.2; 3 — MonoR.3; 4 — PolyG.1; 5 — PolyG.2; 6 — PolyG.3; 7 — PolyC.1; 8 — PolyC.2; 9 — PolyC.3;

**S1 Table. Geobotanical description of the sampling sites**.

## Notes

### Competing Interest Statement

The authors have declared no competing interest.

